# HiCLift: A fast and efficient tool for converting chromatin interaction data between genome assemblies

**DOI:** 10.1101/2023.01.17.524475

**Authors:** Xiaotao Wang, Feng Yue

## Abstract

**Motivation:** With the continuous effort to improve the quality of human reference genome and the generation of more and more personal genomes, the conversion of genomic coordinates between genome assemblies is critical in many integrative and comparative studies. While tools have been developed for such task for linear genome signals such as ChIP-Seq, no tool exists to convert genome assemblies for chromatin interaction data, despite the importance of three-dimensional (3D) genome organization in gene regulation and disease.

**Results:** Here, we present HiCLift, a fast and efficient tool that can convert the genomic coordinates of chromatin contacts such as Hi-C and Micro-C from one assembly to another, including the latest T2T genome. Comparing with the strategy of directly re-mapping raw reads to a different genome, HiCLift runs on average 42 times faster (hours vs. days), while outputs nearly identical contact matrices. More importantly, as HiCLift does not need to re-map the raw reads, it can directly convert human patient sample data, where the raw sequencing reads are sometimes hard to acquire or not available.

**Availability:** HiCLift is publicly available at https://github.com/XiaoTaoWang/HiCLift.

## 1 Introduction

Reference genome is the core of a modern genomic study, which serves as a standardized coordinate system for mapping high-throughput sequencing reads and annotating genomic elements. However, the reference genome of human, mouse, and other species have been and are still being modified over the years. For example, the first draft of human genome published in 2001 contains ~150,000 gaps (Lander, et al., 2001); the GRCh38 version released by the Genome Reference Consortium in 2013 contains 827 gaps (~151 mega-base pairs); and the most recent T2T-CHM13 version released by the Telomere-to-Telomere (T2T) Consortium is a complete sequence of a human genome, which includes gapless assemblies of all chromosomes except Y (Nurk, et al., 2022).

Hundreds of thousands of genomic studies have been performed, and the sequencing data have been analyzed against different versions of reference genome. To facilitate an integrative analysis of the published data, it would be necessary to convert the different sequencing data into a consistent coordinate system. In general, two approaches can be considered. The first approach is to re-map the original sequencing reads to the same target assembly. This approach provides the most accurate result but is computationally intensive and time-consuming. The second approach is to convert the genomic coordinates between assemblies by using a mapping file. Although there can be information loss during the conversion, this approach gives a good trade-off between performance and accuracy for most applications. Several tools, such as UCSC liftOver, NCBI Remap, CrossMap, and segment_liftover, have been developed to perform the coordinate conversion for datasets coming from various genomic/epigenomic experiments (Gao, et al., 2018; Kuhn, et al., 2013; Zhao, et al., 2014). However, despite the importance of 3D genome organization in gene regulation and disease, no tool exists to convert genome assemblies for chromatin interaction data.

Recent years have seen a growth spurt of 3D genome technologies, such as Hi-C (Lieberman-Aiden, et al., 2009), Micro-C (Hsieh, et al., 2015), ChIA-PET (Fullwood, et al., 2009), HiChIP (Mumbach, et al., 2016), GAM (Beagrie, et al., 2017), DNA SPRITE (Quinodoz, et al., 2018), and ChIA-Drop (Zheng, et al., 2019), that map spatial associations between two or more genomic loci. There have been over 600 datasets generated to study the higher-order chromatin structure and the interactions between genes and their distal regulatory elements (Reiff, et al., 2022). In general, each experiment has several hundred million or billions of raw reads. As a matter of fact, a recent study even generated > 7 billion reads in mouse ES cells (Bonev, et al., 2017). Due to deep sequencing, re-processing these data to a specific genome assembly could be extremely time-consuming. More importantly, there have been many Hi-C datasets that were generated for human individuals or patient samples, and the raw sequencing reads are not publicly available, which severely hinder our ability to integrate such data with the newly generated data.

Here, we introduce HiCLift, an efficient command-line tool that can convert genomic coordinates of chromatin contacts between assemblies. HiCLift supports a variety of data formats that are widely used by the 3D genome community, such as 4DN pairs (https://github.com/4dn-dcic/pairix/blob/master/pairs_format_specification.md), allValidPairs outputted by HiC-Pro (Servant, et al., 2015), .cool (Abdennur and Mirny, 2020), and .hic (Durand, et al., 2016). Using large Hi-C datasets of human, mouse, and zebrafish as a benchmark, we show that compared with the strategy directly remapping raw reads to a different genome, HiCLift runs on average ~42 times faster, while outputs nearly identical contact matrices.

## 2 Materials and Methods

### 2.1 Hi-C data sources and processing

The Hi-C datasets of a human fibroblast cell line IMR90 and a mouse lymphoma cell line CH12-LX were downloaded from the GEO database with accession code GSE63525 (Rao, et al., 2014). The Hi-C dataset of a zebrafish muscle tissue was downloaded from GEO with accession code GSE134055 (Yang, et al., 2020). The Micro-C dataset of H1-ESC cells were downloaded from GEO with accession code GSE163666 (Akgol Oksuz, et al., 2021). The raw sequencing reads were processed using runHiC (https://pypi.org/project/runHiC/, v0.8.6), a command-line tool based on the 4D Nucleome Hi-C data processing pipeline (https://data.4dnucleome.org/resources/data-analysis/hi_c-processing-pipeline). To benchmark the performance of HiCLift, each dataset was first processed into two versions of chromatin contacts using two different genome assemblies, and then the coordinates of chromatin contacts mapped to the older genome version were converted into the newer genome version using HiCLift, and compared with chromatin contacts directly mapped the newer genome version. Specifically, the IMR90 dataset was processed into hg38 and T2T-CHM13 (v2.0), the CH12-LX dataset was processed into mm10 and mm39, and the zebrafish muscle dataset was processed into danRer10 and danRer11.

We compared the contact maps derived from runHiC and HiCLift at different scales. First, we measured the overall similarity between two contact maps by using the stratum-adjusted correlation coefficients (SCC) at the 50kb resolution (Yang, et al., 2017). Second, we compared the chromatin compartments measured by the first eigenvector (PC1) of the normalized contact matrices at 100kb. Third, we compared the domain boundary strength measured by the insulation scores at 25kb. Finally, we compared the identified chromatin loops at the 5kb resolution.

Specifically, SCCs were computed using a Python implementation of the original HiCRep algorithm (https://github.com/dejunlin/hicrep, v0.2.6), with the smoothing factor and the maximum genomic distance set to 3 and 5Mb, respectively. The weighted average of SCCs from individual chromosomes (using chromosome lengths as the weights) were reported as the final SCC score for each comparison. Both compartments and TADs were estimated using cooltools (https://pypi.org/project/cooltools/, v0.4.0). For compartments, the eigenvalue decomposition was performed on the 100kb intra-chromosomal contact maps, and the first eigenvector (PC1) was used to capture the “plaid” contact pattern. The original PC1 was oriented according to H3K4me3 ChIP-Seq tracks of corresponding cells, so that positive values correspond to active genomic regions and negative values correspond to inactive regions. For TADs, genome-wide insulation scores (IS) were calculated at 25kb with the window size setting to 500kb. Finally, the chromatin loops were identified at 5kb resolution using a Python implementation of the HiCCUPS algorithm (https://pypi.org/project/hicpeaks/, v0.3.5) (Salameh, et al., 2020).

### 2.2 ChlP-Seq data sources and processing

As mentioned above, we used H3K4me3 ChIP-Seq tracks to orient the original PC1 so that the positive PC1 values correspond to active regions and negative PC1 values correspond to inactive regions. For IMR90, we downloaded the track from ENCODE (https://www.encodeproject.org/) with accession code ENCFF518GFI, and converted the original coordinates from hg38 to T2T-CHM13 using CrossMap (v0.5.2) (Zhao, et al., 2014). For CH12-LX, we downloaded the track from ENCODE with accession code ENCFF012DBS, and converted the coordinates from mm10 to mm39. For zebrafish muscle, we downloaded the raw sequencing reads from the SRA database with accession code SRR9662073, mapped the reads to danRer11 using BWA-MEM (v0.7.17), and generated the signal track using “macs2 callpeak” (v2.2.7.1) with “-B” and “--SPMR” parameters.

### 2.3 Overview of HiCLift

The inputs to HiCLift include two parts (Fig. 1). The first part is a file containing the chromatin contact information. This file can be either a pairs file (4DN pairs or HiC-Pro allValidPairs) with each row representing a pair of interacting genomic loci in base-pair resolution, or a matrix file (.cool or .hic), which stores interaction frequencies between genomic intervals of fixed size. The second part is a UCSC chain file (https://genome.ucsc.edu/goldenPath/help/chain.html), which describes pairwise alignment that allows gaps in both assemblies simultaneously. Internally, HiCLift represents a chain file as IntervalTrees, with one tree per chromosome, to efficiently search for a specific genomic position in a chain file and locate the matched position in the target genome. The converted chromatin contacts will be reported in either a sorted 4DN pairs file, which can be directly used to generate contact matrix in various formats, or a matrix file in .cool or .hic formats.

**Fig. 1.**
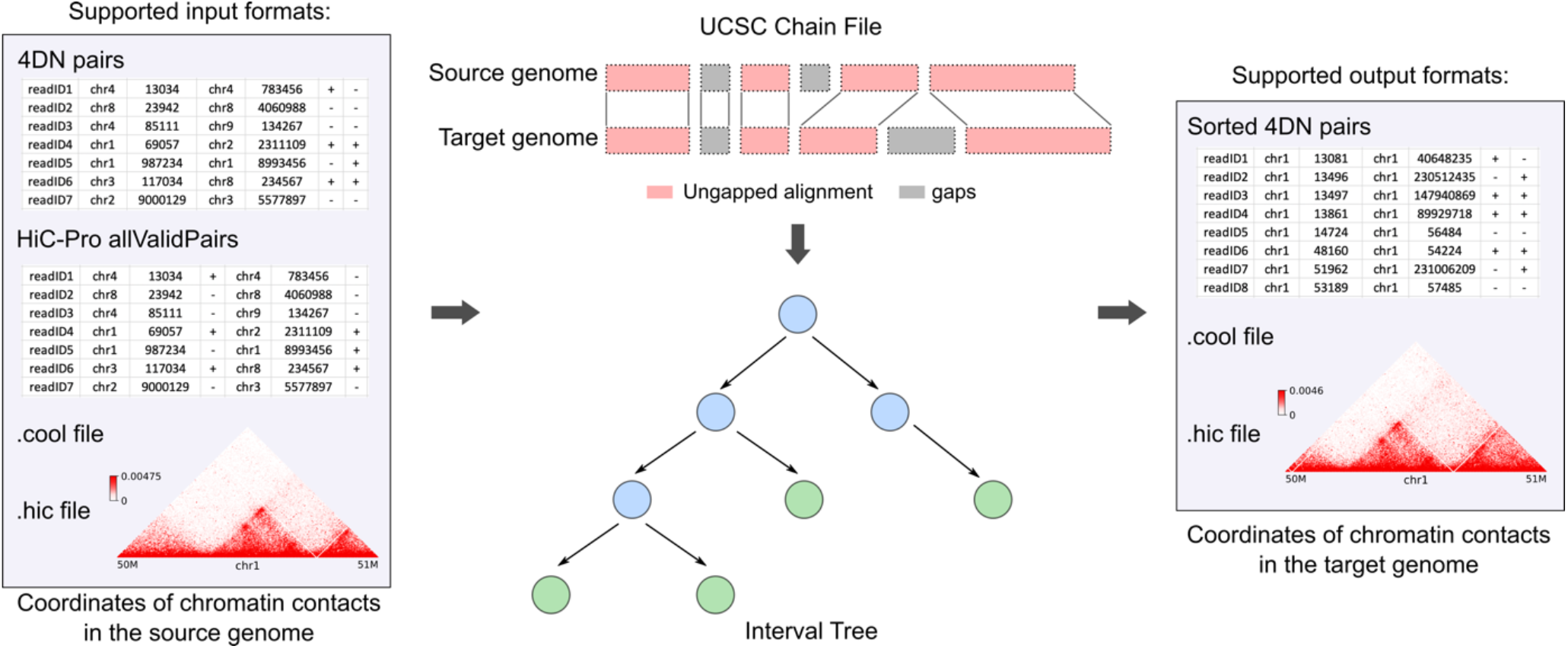
The overall design of HiCLift.

### 2.4 Processing contact pairs

HiCLift supports two kind of pairs files: the pairs format defined by the 4D Nucleome Data Coordination and Integration Center (DCIC) (https://github.com/4dn-dcic/pairix/blob/master/pairs_format_specification.md) and allValidPairs defined by HiC-Pro (https://nservant.github.io/HiC-Pro/RESULTS.html). Both formats define contact pairs in base-pair resolution, with each row representing genomic coordinates of a pair of interacting genomic loci. HiCLift iterates each row of a pairs file, searches for the coordinates in the IntervalTree constructed from the input chain file, and maps to the target genome. A pair of loci is retained only if both sides can be uniquely mapped to the target genome. The input pairs file can be plain text file, gzip/bgzip compressed file (.gz) or lz4 compressed file (.lz4).

### 2.5 Processing contact matrices

Suppose there is a contact matrix ***M***, where each value in the matrix *M_ij_* represents the contact frequency/count between bin *i* and bin *j*. At a given resolution, each bin represents a genomic interval of fixed size. Therefore, the precise interacting loci for each contact are unknown in such a matrix. To maximize the mappability ratio, for each pair of bins, HiCLift searches for loci that can be uniquely mapped to the target genome, and randomly samples a pair of mappable loci for each contact between corresponding bins.

Two matrix formats are supported: cool (Abdennur and Mirny, 2020) and hic (Durand, et al., 2016). Both are official data formats for the 4D Nucleome consortium. For the hic format, since multiple matrices at various resolutions are stored in a single file, HiCLift automatically detects and reads data from the one at the highest resolution.

### 2.6 Source of chain files

The chain file “grch38-chm13v2.chain” for mapping coordinates from hg38 to T2T-CHM13 was downloaded from https://github.com/marbl/CHM13. Other chain files used in this study were downloaded from https://hgdownload.soe.ucsc.edu/downloads.html.

## 3 Results

### 3.1 HiCLift accurately converts genomic coordinates of chromatin contacts from one assembly to another

We benchmarked the performance of HiCLift using three Hi-C datasets of different species. Among them, the IMR90 dataset contains ~1.54 billion read pairs, with ~0.99 billion pairwise contacts after mapping reads to the hg38 human genome assembly and removing PCR duplicates; the CH12-LX dataset contains ~1.38 billion reads and ~0.71 billion pairwise contacts (with reads mapped to the mm10 mouse genome); and the zebrafish muscle dataset contains ~1.35 billion reads and ~0.57 billion contacts (with reads mapped to the danRer10 genome). Using HiCLift, we converted the coordinates of contact pairs (in the 4DN pairs format) for IMR90, CH12-LX, and zebrafish muscle into the most recent T2T-CHM13 human genome assembly (v2.0) (Nurk, et al., 2022), the mm39 mouse genome, and the danRer11 zebrafish genome, respectively. At the same time, we processed these datasets by remapping reads to the same target genome using a regular Hi-C data processing pipeline, and used the result contact maps as a reference to evaluate the accuracy of HiCLift.

We first compared the overall distribution of chromatin contacts at 50kb resolution by using the stratum-adjusted correlation coefficients (SCC) (Yang, et al., 2017). The SCCs between HiCLift and read re-mapping achieved to 0.9987, 0.9997, and 0.9883 for IMR90, CH12-LX, and zebrafish muscle, respectively, which suggests that the coordinates converted from other genome assembly versions by HiCLift are highly concordant with the coordinates from read re-mapping.

We further evaluated the concordance of HiCLift and read re-mapping in detecting downstream chromatin contact features, such as chromatin compartments, topologically associating domains (TADs), and chromatin loops (Fig. 2 and Supplementary Figs. S1-S2). Both chromatin compartments (measured by PC1) and TADs (measured by insulation scores) called from HiCLift and read remapping gave highly similar results, achieving a coefficient of determination (*R*^2^) of 0.9995 and 0.9905 respectively for the IMR90 dataset, 1.0000 and 0.9957 for the CH12-LX dataset, and 0.9995 and 0.9635 for the zebrafish muscle dataset. Using HiCCUPS (Rao, et al., 2014), we detected 8,258 chromatin loops for the HiCLift-converted version of the IMR90 contact map at 5kb resolution, 94.4% (7,799/8,258) of which exactly matched 93.9% (7,799/8,310) of loops detected from the read remapping version at the same resolution. Similarly, for the CH12-LX dataset, 99.2% (7,778/7837) of loops from HiCLift matched 98.9% (7,778/7,867) of loops from read re-mapping; and for the zebrafish muscle dataset, 95.1% of loops from HiCLift overlapped with 89.0% of loops from read re-mapping.

**Fig. 2.**
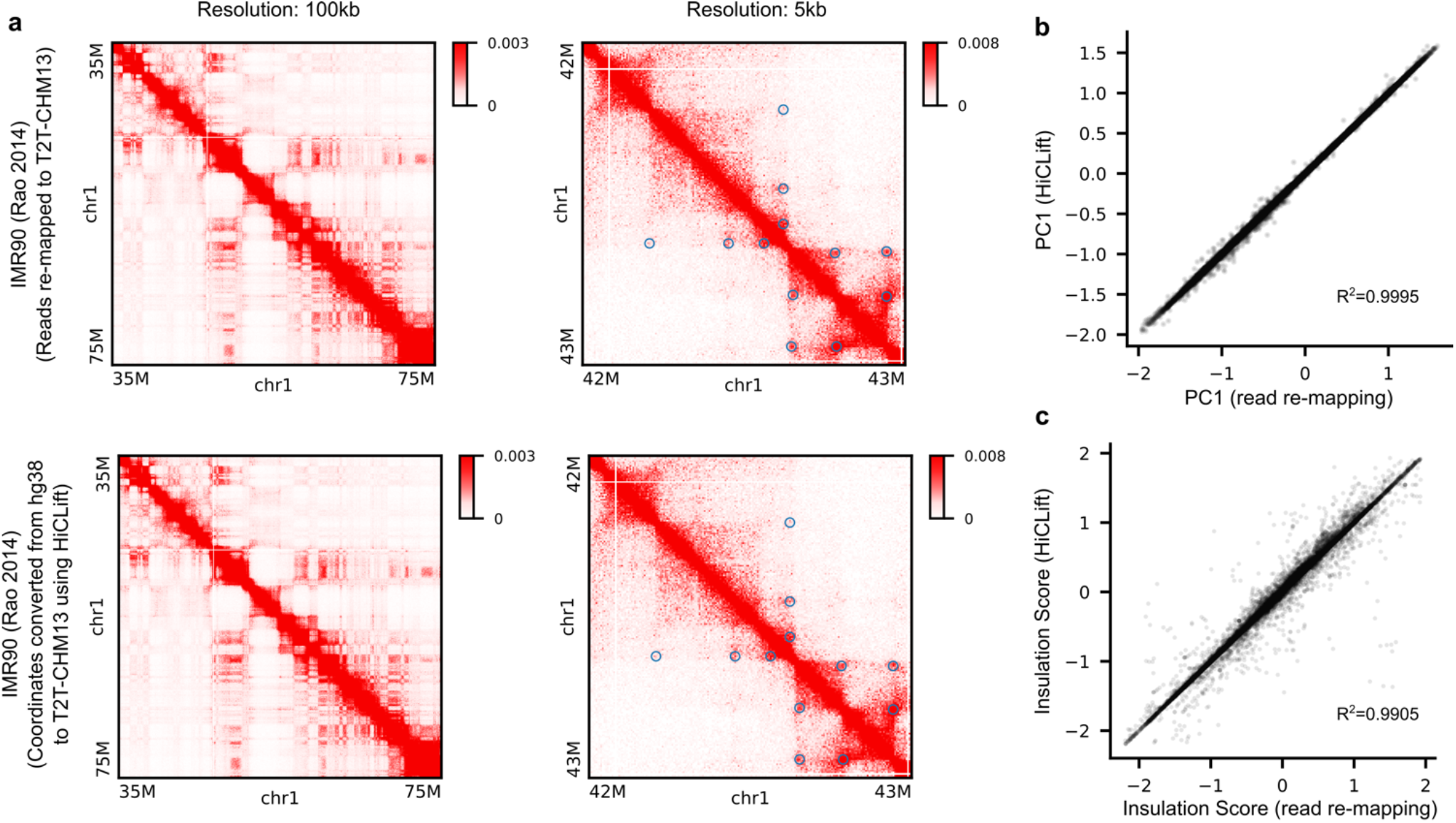
Performance of HiCLift in converting Hi-C data from hg38 to T2T genome. We used the IMR90 Hi-C dataset from Rao 2014, which contains ~1.54 billion raw reads. **(a)** Example regions comparing contact matrices obtained from HiCLift (second row) with the matrices we generated by re-mapping the raw reads (first row). The blue circles in the second column indicate the detected chromatin loops on corresponding maps. **(b)** The first principal component (PC1) for characterizing the chromatin compartment pattern at 100kb resolution is compared between the two methods. **(c)** The insulation scores for capturing chromatin domain boundaries at 25kb resolution are compared between the two methods.

In addition to contact pairs at the base-pair resolution, we also evaluated the accuracy of HiCLift when contact matrices binned at coarser resolutions are used as input. As expected, the SCCs between HiCLift and read re-mapping decreased as the bin sizes of the input matrix increased (Supplementary Figs. S3-S5). However, even with the 10kb contact matrices as input, the SCCs still achieved 0.9986, 0.9992, and 0.9853 for the IMR90, CH12-LX, and zebrafish muscle datasets, respectively.

Together, these analyses show that HiCLift can accurately convert genomic coordinates of chromatin contacts between different genome assemblies for various species.

### 3.2 HiCLift accurately converts contact coordinates at erroneously assembled regions

Older reference genome versions usually contain higher number of erroneously assembled regions than newer versions. To investigate whether HiCLift works for these regions, we focused on the human *H19/ICF2* locus, where the *LINC01150* gene at this locus has been shown to be inverted and translocated upstream of the *TNNT3* gene in the hg38 genome assembly, compared with the most recent T2T-CHM13 genome (Battaglia, et al., 2022) (Fig. 3a). Consistent with this, we found when reads from a Micro-C dataset were mapped to hg38, abnormal interaction blocks showed up between *LINC01150* and *TNNT3;* however, when reads were mapped to T2T-CHM13, no such blocks could be observed (Fig. 3b). And consistent with our previous evaluation, lifting over contact coordinates from hg38 to T2T-CHM13 with HiCLift produced nearly identical contact matrices to the one with reads directly mapped to T2T-CHM13.

**Fig. 3.**
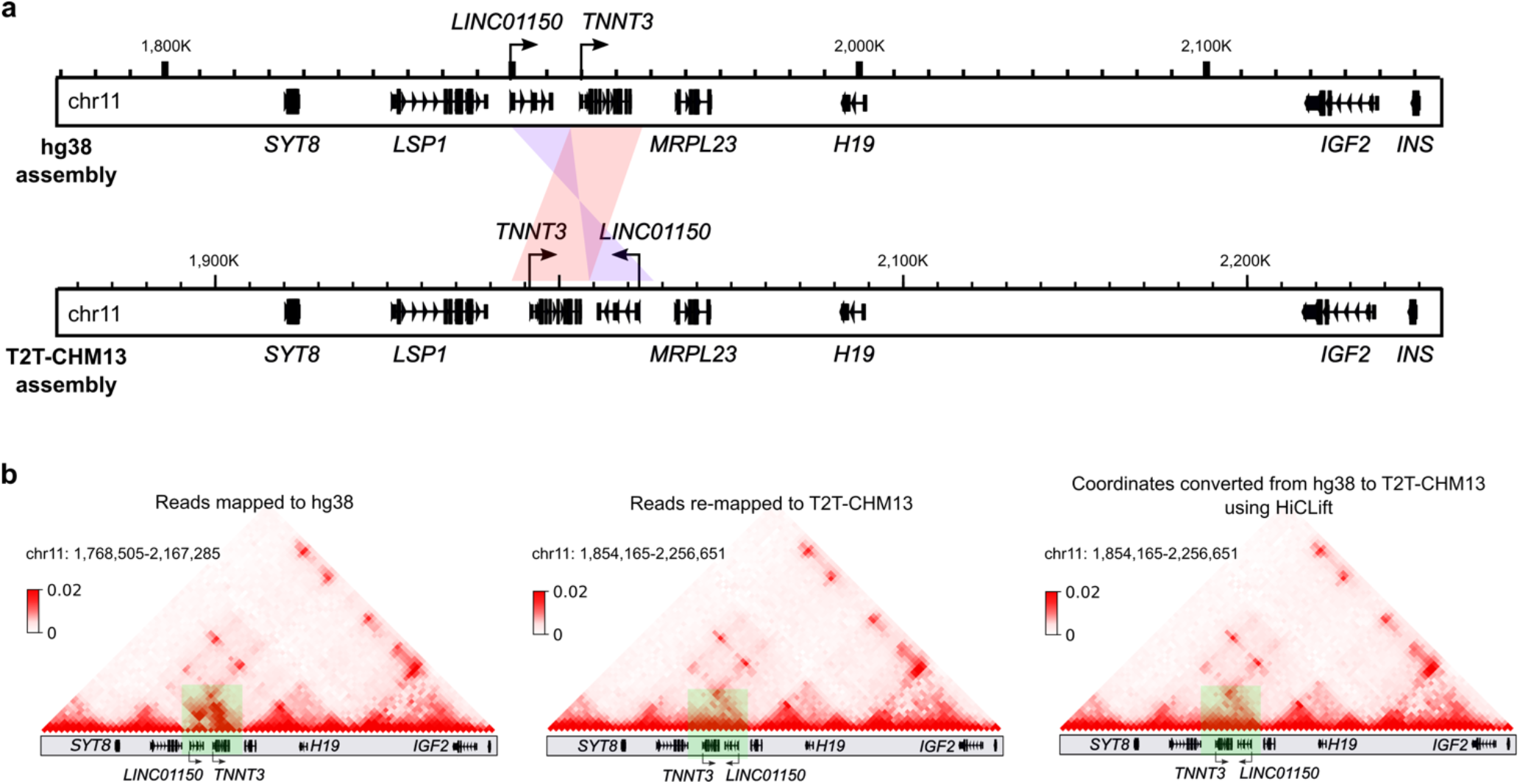
Accuracy of HiCLift at erroneously assembled regions. **(a)** Comparisons of hg38 and T2T-CHM13 genomes show that the *LINC01150* gene is inverted and inserted upstream of *TNNT3* in the hg38 assembly. **(b)**. Comparisons of chromatin contacts (from a Micro-C dataset in H1-ESC cells) mapped to hg38, mapped to T2T-CHM13, and converted from hg38 to T2T-CHM13. The highlighted regions show that abnormal chromatin contacts exist in hg38 due to assembly errors, but are absent from T2T-CHM13.

Similarly, disease-associated regions at 16p12.1 were reported to be misoriented in hg38 (Antonacci, et al., 2010). When we examined the H1-ESC Micro-C data mapped to hg38, there were many aberrant off-diagonal signals. However, such signals were no longer visible on the map either with reads remapped to T2T-CHM13 or with coordinates converted from hg38 to T2T-CHM13 (Supplementary Fig. S6).

Together, HiCLift can accurately convert contact coordinates, even at erroneously assembled regions.

### 3.3 HiCLift runs 42 times faster than read re-mapping

As 3D genome libraries usually need deep sequencing for effective downstream analysis, processing these data from scratch is often time-consuming. For example, processing the benchmark datasets in this study from raw sequencing reads to sorted 4DN pairs took ~52.8 hours, 45.2 hours, and 84.9 hours for IMR90, CH12-LX, and zebrafish muscle, respectively (8 CPU cores were allocated). As a comparison, HiCLift (8 CPU cores were allocated) took ~2.2 hours (24x), 1.4 hours (32x), and 1.2 hours (71x) to finish the coordinate conversion, which was on average 42 times faster than a standard Hi-C data processing pipeline (Fig. 4a). To test the performance of HiCLift at various sequencing depths, we computationally down-sampled the IMR90 dataset into 9 different depths (ranging from 100 million to 900 million contact pairs). Overall, the running time for HiCLift grew linearly with the sequencing depths. Notably, datasets with fewer than 500 million contact pairs only needed <1 hour for HiCLift to finish the conversion (Fig. 4b).

**Fig. 4.**
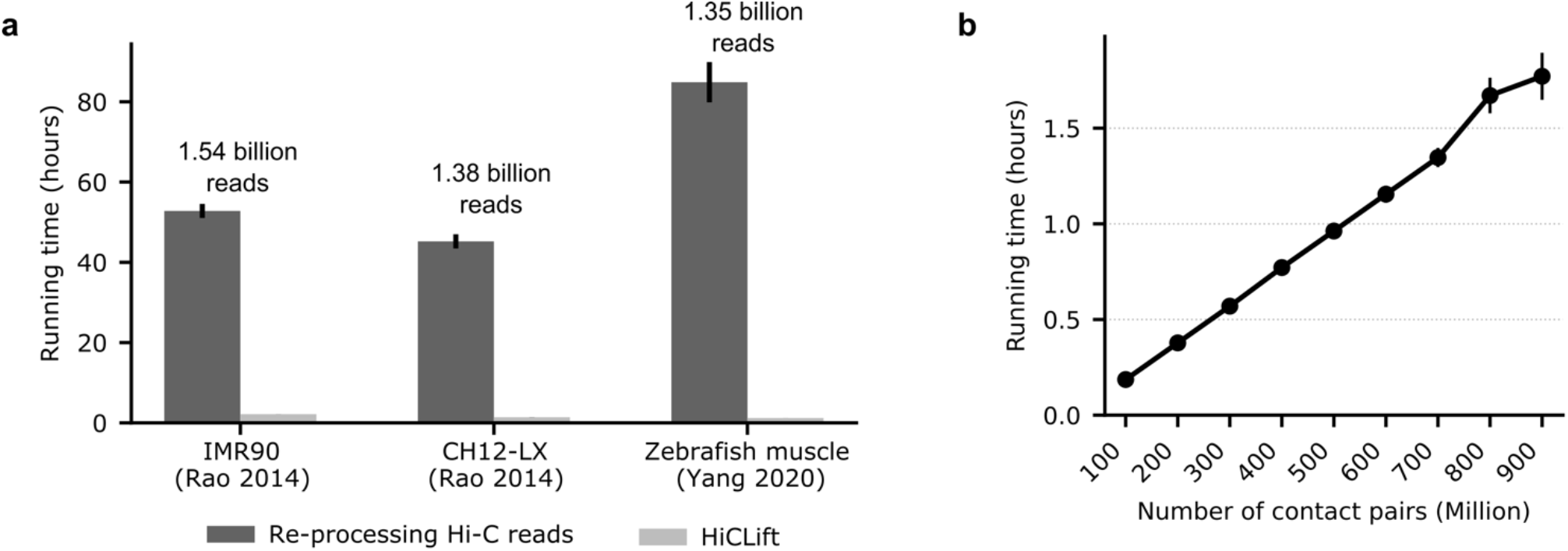
Computational efficiency of HiCLift on the benchmark datasets. **(a)** Running time comparison of HiCLift with Hi-C data re-processing on the benchmark datasets. For each dataset and each method, we repeated the computation for 3 times, and the error bars represent the standard deviation for corresponding 3 repeats. **(b)** Running time of HiCLift on the down-sampled IMR90 datasets. The error bars represent the standard deviation for 5 repeats at each sequencing depth.

## 4 Discussion

In this work, we developed a novel computational tool HiCLift and demonstrated its accuracy and efficiency for converting genomic coordinates of chromatin contacts between assemblies. Although one may apply the tool with arbitrary chain files, like other liftover tools, HiCLift was optimized only for intraspecies genomic conversions. As the volume of chromatin interaction data keeps increasing, we envision HiCLift to play a vital role in an integrative or comparative analysis when re-mapping raw reads is not feasible.

---

**Supplementary Fig. S1.**
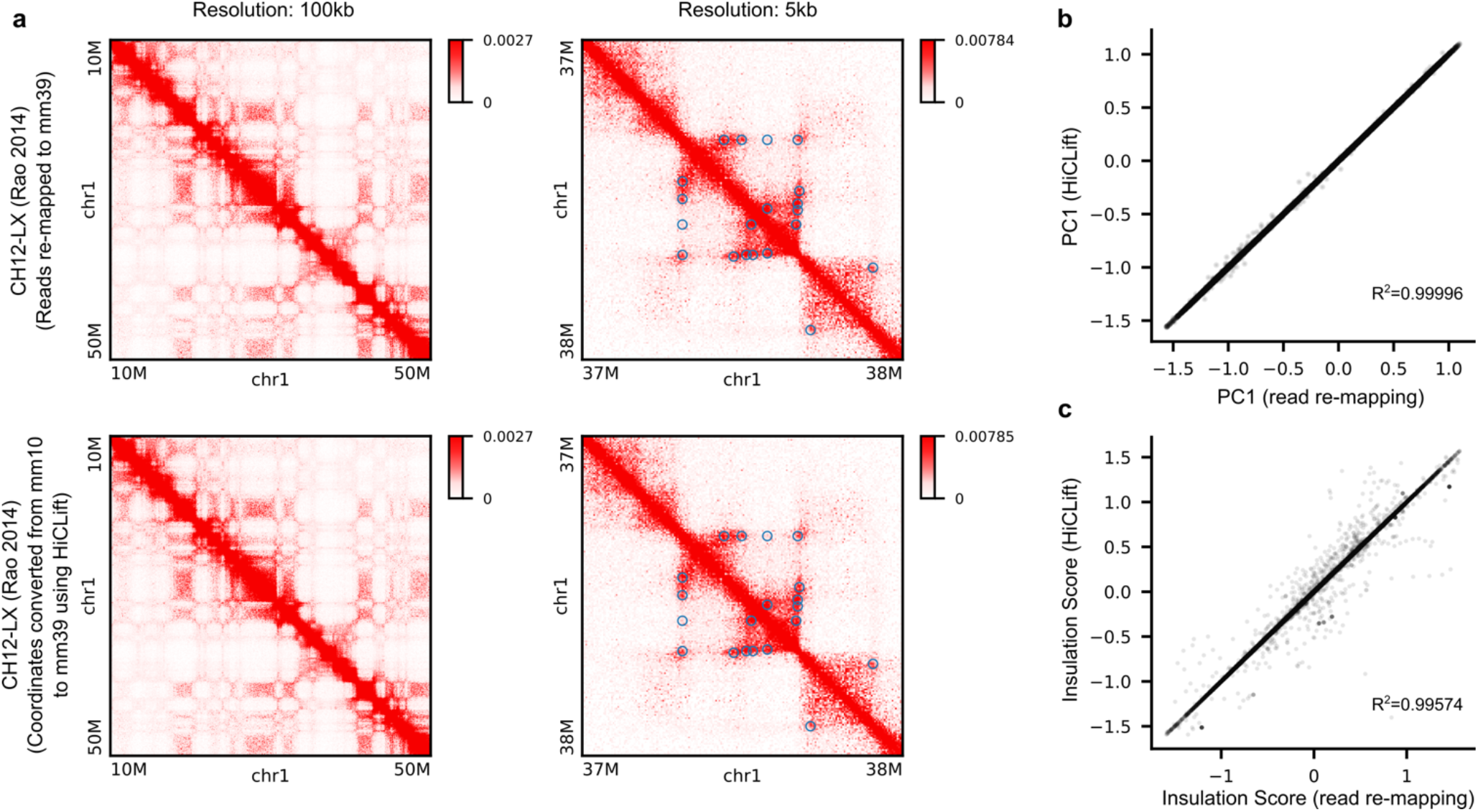
Accuracy of HiCLift on a mouse Hi-C dataset (Rao 2014, CH12-LX). **(a)** Example regions comparing contact matrices obtained from HiCLift with the matrices from read remapping. The blue circles indicate the detected chromatin loops on corresponding maps. **(b)** The first principal component (PC1) for characterizing the chromatin compartment pattern at 100kb resolution is compared between the two methods. **(c)** The insulation scores for capturing chromatin domain boundaries at 25kb resolution are compared between the two methods.

**Supplementary Fig. S2.**
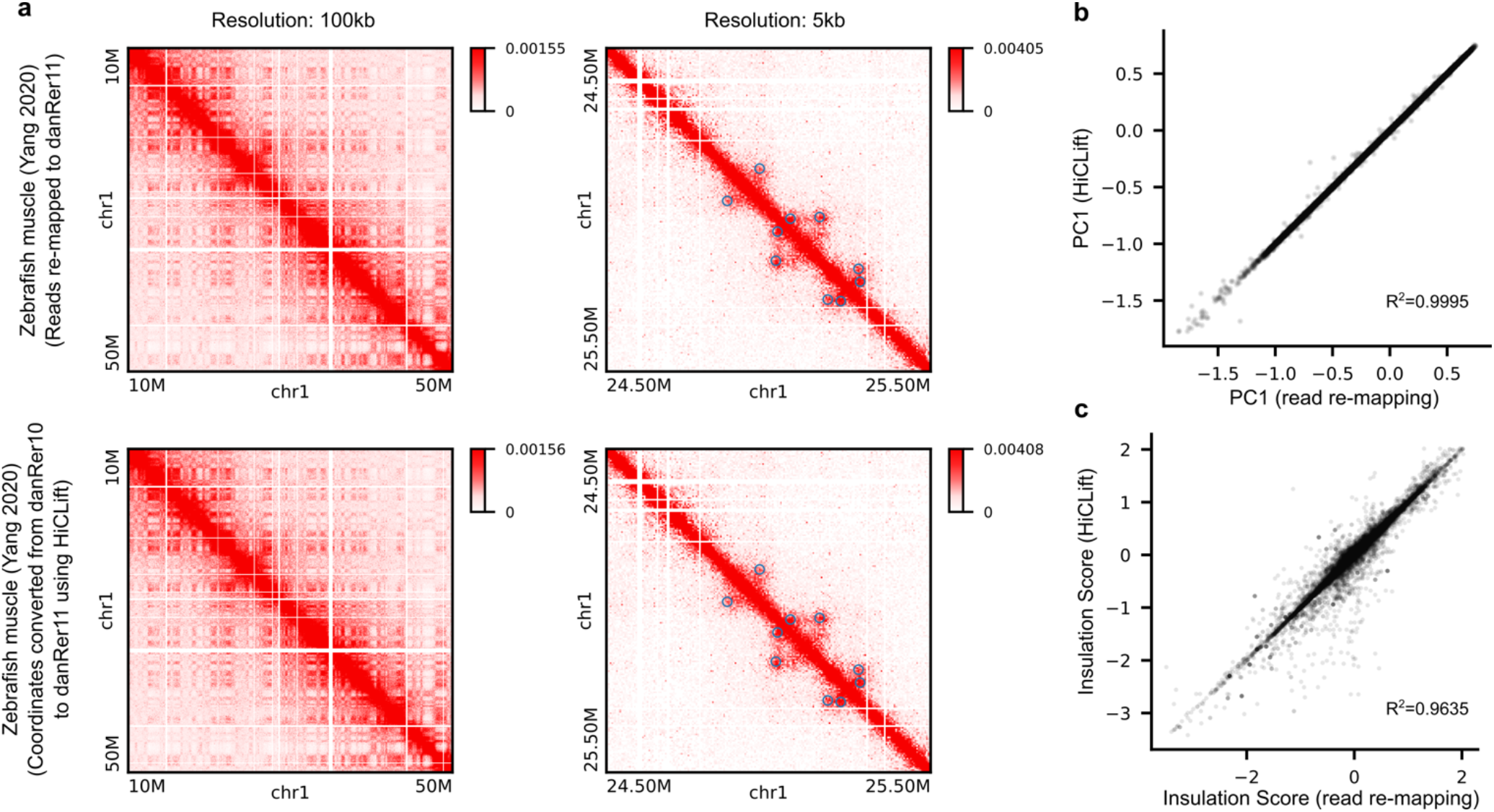
Accuracy of HiCLift on a zebrafish Hi-C dataset (Yang 2020, muscle tissue). **(a)** Example regions comparing contact matrices obtained from HiCLift with the matrices from read re-mapping. The blue circles indicate the detected chromatin loops on corresponding maps. **(b)** The first principal component (PC1) for characterizing the chromatin compartment pattern at 100kb resolution is compared between the two methods. **(c)** The insulation scores for capturing chromatin domain boundaries at 25kb resolution are compared between the two methods.

**Supplementary Fig. S3.**
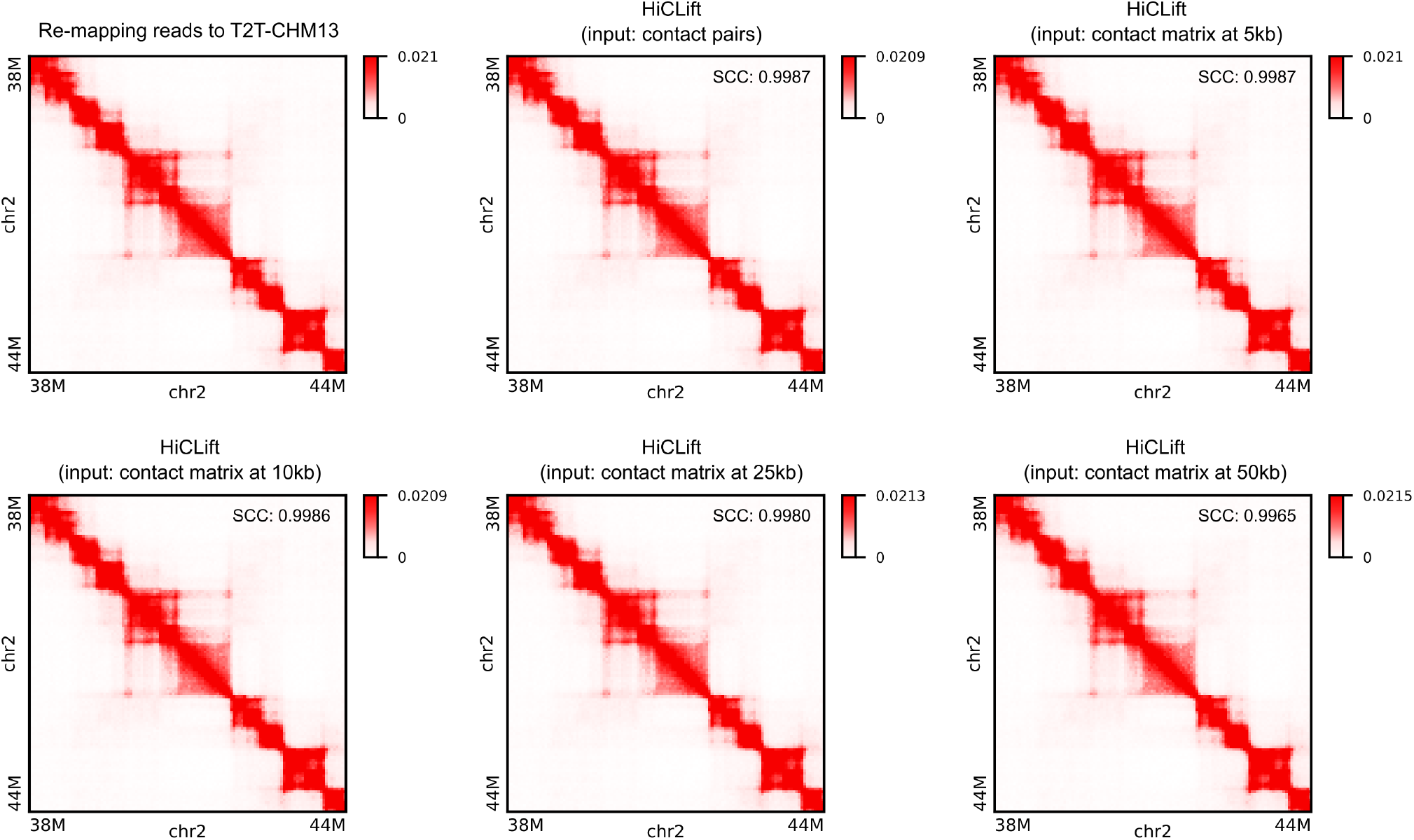
Comparison of Hi-C contact maps at 50kb resolution derived from read re-mapping and HiCLift on the IMR90 dataset. We ran HiCLift with contact pairs, and contact matrices at different resolutions as input to convert contact coordinates from hg38 to T2T-CHM13. The results are highly similar to the one with reads re-mapped to T2T-CHM13 regardless of the input data format and resolution. The stratum-adjusted correlation coefficient (SCC) between HiCLift and read re-mapping is indicated in each case.

**Supplementary Fig. S4.**
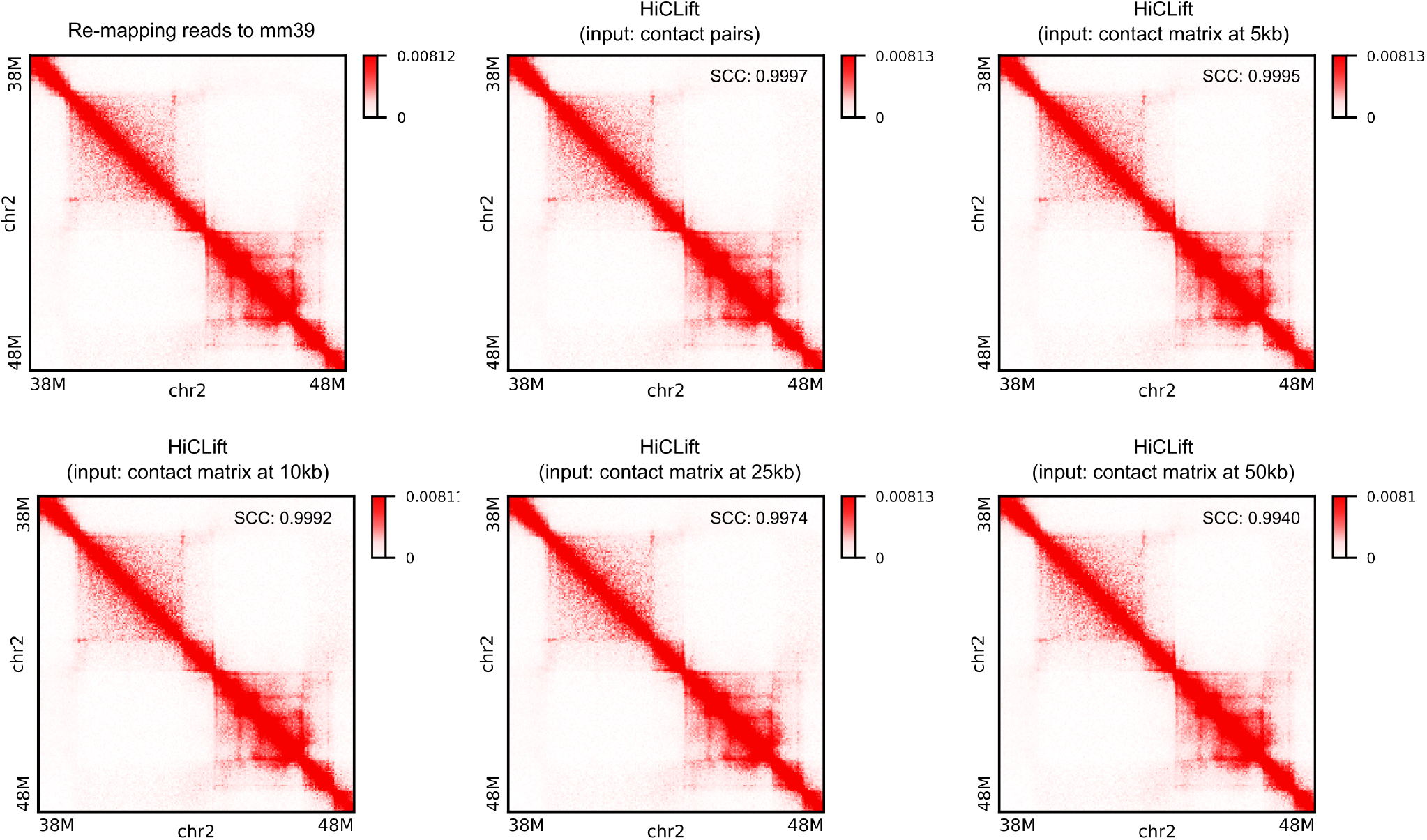
Comparison of Hi-C contact maps at 50kb resolution derived from read remapping and HiCLift on the CH12-LX dataset. We ran HiCLift with contact pairs, and contact matrices at different resolutions as input to convert contact coordinates from mm10 to mm39. The results are highly similar to the one with reads re-mapped to mm39 regardless of the input data format and resolution. The stratum-adjusted correlation coefficient (SCC) between HiCLift and read re-mapping is indicated in each case.

**Supplementary Fig. S5.**
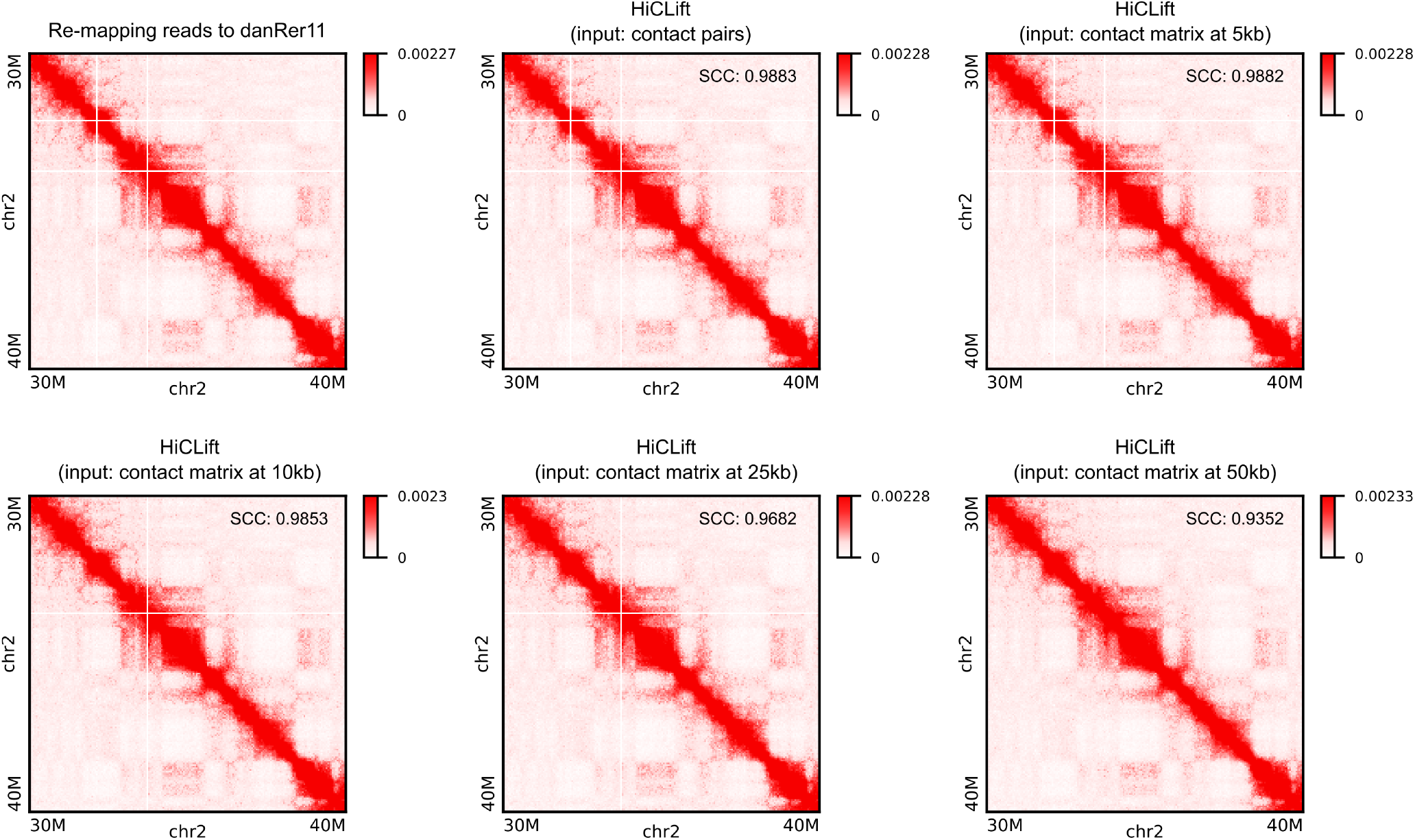
Comparison of Hi-C contact maps at 50kb resolution derived from read remapping and HiCLift on the zebrafish dataset. We ran HiCLift with contact pairs, and contact matrices at different resolutions as input to convert contact coordinates from danRer10 to danRer11. The results are highly similar to the one with reads re-mapped to danRer11 regardless of the input data format and resolution. The stratum-adjusted correlation coefficient (SCC) between HiCLift and read re-mapping is indicated in each case.

**Supplementary Fig. S6.**
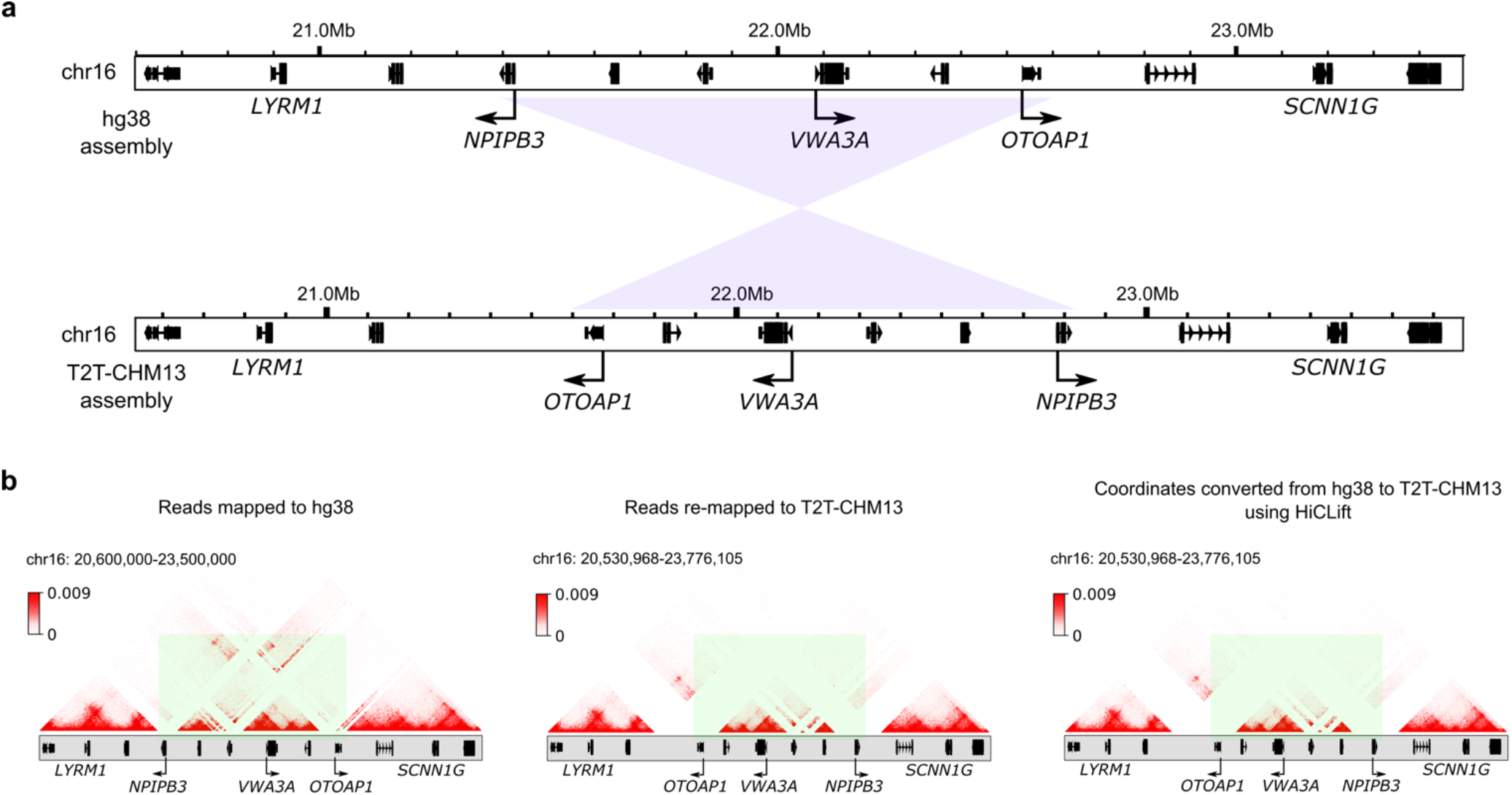
Accuracy of HiCLift at erroneously assembled regions. **(a)** Comparisons of hg38 and T2T-CHM13 show that a 1Mb-fragment encompassing the genes *NPIPB3*, *VWA3A*, and *OTOAP1* is inverted in hg38. **(b)**Comparisons of chromatin contacts (from a Micro-C dataset in H1-ESC cells) mapped to hg38, mapped to T2T-CHM13, and converted from hg38 to T2T-CHM13. The highlighted regions show that abnormal chromatin contacts exist in hg38 due to assembly errors, but are absent from T2T-CHM13.

